# Muscarinic acetylcholine receptor activated effectors in principal neurons of the rat basolateral amygdala

**DOI:** 10.1101/2025.03.14.643393

**Authors:** Todd J. Sahagian, Scott W. Harden, Jennifer L. Bizon, Barry Setlow, Charles J. Frazier

## Abstract

The basolateral amygdala (BLA) plays a crucial role in context-specific learning and memory by integrating valence-specific stimuli with internal physiological states. Cholinergic signaling systems modulate neural excitability to influence information processing in the BLA. Muscarinic acetylcholine receptors (mAChRs) are of particular interest because aberrant mAChR signaling in BLA circuits is associated with neuropsychiatric disorders, cognitive impairment, substance use, and age-related cognitive decline. This study evaluates mAChR activation in BLA principal neurons (PNs) in juvenile rat brain slices using whole-cell patch-clamp recordings. We found that bath application of carbachol (CCh,) produces a pirenzepine sensitive excitatory response in BLA PNs voltage clamped near the resting potential, which depends on an underlying biphasic change in membrane resistance, indicating an involvement of multiple effectors. More specifically, we observed that CCh excites BLA PNs by inhibiting the afterhyperpolarization (AHP), by reducing a steady state inhibitory current, and by promoting an afterdepolarization (ADP). We further identify and characterize a CCh-induced and calcium-activated non-selective cation current (I_CAN_) that underlies the ADP in voltage clamp. Overall, our findings provide new insights into specific effectors modulated by activation of pirenzepine sensitive mAChRs expressed by BLA PNs. We also reveal new details about the time-and voltage-dependence of current carried by the CCh - activated I_CAN_ like current in BLA PNs, and highlight its ability to promote a suprathreshold ADP capable of generating sustained firing after a brief excitatory stimulus. Improved understanding of these effectors will provide potentially valuable new insights on the wide range of mechanisms through which cholinergic system dysfunction can lead to impaired executive function.

## 1. Introduction

Neural processing of information traveling through BLA circuitry impacts the balance of fear, stress, motivation, decision-making, and reward processing, which underlie complex behaviors that facilitate survival. Aberrant signaling in the BLA is associated with clinical manifestations such as post-traumatic stress disorder, substance use disorders, anxiety disorders, aggression, and impulsivity, but despite its important role in health and disease the mechanisms that mediate information processing through the BLA remain incompletely understood [1–5]. The BLA receives heterogeneous synaptic inputs from numerous brain centers including the lateral amygdala, hippocampus, prefrontal cortex, and the bed nucleus of the stria terminals, and the BLA also sends projections to many locations including the central amygdala, medial amygdala, prefrontal cortex, thalamus, hypothalamus, and nucleus accumbens [6]. Glutamatergic inputs to the BLA are the primary source of fast excitatory neurotransmission, but BLA neurons are sensitive to a wide variety of neurotransmitters and neuromodulators including acetylcholine, norepinephrine, serotonin, cholecystokinin, corticotropin releasing hormone, endocannabinoids, and more, which modify their excitability to fine-tune how information is processed as it travels through the BLA [7–11].

Cholinergic signaling and its ability to modulate information flow through the BLA is of particular interest because pharmacological interventions that target acetylcholine receptors (AChRs) have the potential to improve anxiety, schizophrenia, mood disorders, substance use, and age-related cognitive decline [12–17]. The BLA has a high density of cholinergic terminals originating from the basal forebrain and brainstem. BLA PNs express muscarinic acetylcholine receptors (mAChRs) which can be studied experimentally using carbachol (CCh), a non-selective cholinergic agonist [18]. CCh sensitive mAChRs expressed by BLA PNs ultimately activate several distinct effector systems [19,20]. A major excitatory action of CCh on BLA PNs is caused by the closing of potassium channels that are open near the resting membrane potential, which depolarizes neurons, and also increases their sensitivity to excitatory synaptic inputs due to increased membrane resistance. Activation of mAChRs also influences complex firing behavior of BLA PNs. Strong firing activates a milieu of voltage-gated and/or calcium-activated currents to produce an inhibitory afterhyperpolarization (AHP) and an excitatory afterdepolarization (ADP), which combine in complex ways to influence synaptic integration by modulating the probability and timing of subsequent action potentials. Unal et al. [21] demonstrated that optogenetic release of ACh in the BLA results in a large activity-dependent ADP that is sensitive to flufenamic acid (FFA), suggesting it may be caused by a calcium-activated nonselective cation current (I_CAN_). Activation of mAChRs also reduces calcium-activated potassium currents underlying the slow AHP (sI_AHP_), and the voltage-dependent potassium current (I_M_), which collectively further contributes to a net excitatory effect [20,22].

In this study we used whole-cell patch-clamp techniques to further characterize how mAChR activation influences excitability of BLA PNs in acute rat brain slices. Consistent with previous reports, we found that mAChR activation at subthreshold potentials produces a depolarizing current through the combined action of closing potassium channels and opening calcium-dependent and likely nonselective cation channels. In addition, we find that CCh produces a calcium-dependent and FFA-sensitive ADP, and we extend previous reports by isolating this current (I_ADP_) in voltage-clamp configuration to evaluate its time dynamics and voltage dependence. In addition to these findings, the method we describe to isolate and characterize the I_ADP_ can be applied to future studies that probe alterations in cholinergic signaling associated with various physiological and pathological states that influence BLA-dependent behaviors.

## 2. Methods

### 2.1. Acute Brain Slice Preparation

All animal procedures were reviewed and approved by the University of Florida Institutional Animal Care and Use Committee (IACUC). Juvenile male Sprague-Dawley rats (P16-28) were used for this study. Acute brain slices were prepared as previously described [23]. Briefly, rats were administered a ketamine/xylazine cocktail (100 mg/kg ketamine, 10 mg/kg xylazine, i.p.) and after adequate anesthesia was achieved (no response to hind-paw pinch) rats were rapidly decapitated and brains were rapidly extracted and submerged in ice-cold sucrose-laden artificial cerebrospinal solution (ACSF) containing (in mM) 205 sucrose, 10 dextrose, 1 MgSO4, 2 KCl, 1.25 NaH2PO4, 1 CaCl2, and 25 NaHCO3, oxygenated with carbogen (95% O2 / 5% CO2). Brains were coronally sectioned at 300 microns using a Leica VT1200S vibratome, then transferred to an incubation solution containing (in mM) 124 NaCl, 10 dextrose, 3 MgSO_4_, 2.5 KCl, 1.23 NaH_2_PO_4_, 1 CaCl_2_, 25 NaHCO_3_, oxygenated with carbogen and maintained at 35 ºC for 30 minutes. Brain slices were then permitted to passively equilibrate to room temperature for at least 30 minutes prior to recording. Brain slices were transferred to a perfusion chamber and submersed in ACSF containing (in mM) 126 NaCl, 11 dextrose, 1.5 MgSO_4_, 3 KCl, 1.2 NaH_2_PO_4_, 2.4 CaCl_2_, 25 NaHCO_3_, continuously oxygenated with carbogen, perfused at 2 mL/min, and maintained at 28 ºC. ACSF for all experiments was supplemented with glutamate-and GABA receptor antagonists (20 µM DNQX, 40 µM AP5, 100 µM PTX, and 10 µM CGP-55845) to minimize influence of synaptic signaling (on AMPA, NMDA, GABA_A_, and GABA_B_ receptors, respectively) and enhance measurement of isolated somatic currents.

### 2.2. Electrophysiology

Brain slices were visualized using an Olympus BX-WI upright stereomicroscope equipped for infrared differential interference contrast microscopy, a 12-bit CCD camera (QImaging Rolera-XR), and micro-manager software [24]. Patch-clamp experiments were performed using a Multiclamp 700B patch-clamp amplifier with a CV-7B headstage, a DigiData 1440A digitizer (Molecular Devices), and pClamp 11 software. Data were obtained at 20 kHz and voltage-clamp data were filtered using a 7-pole Bessel low-pass filter with a 2 kHz −3 dB cutoff. On-line analysis was performed with custom software written using Python 3.11 and the pyABF package. Off-line analysis was performed with OriginPro (OriginLab, Northampton, MA) using custom software written in OriginC by CJF.

### 2.3. Intracellular Solutions

Patch pipettes were pulled from borosilicate glass (Sutter Instrument BF150-86-10) using a Flaming/Brown pipette puller (Sutter Instrument SU-P97) and had an open-tip resistance of 4-6 MΩ when filled with a K-gluconate based internal solution containing (in mM) 115 K-gluconate, 10 di-tris phosphocreatine, 10 HEPES, 0.5 EGTA, 2 MgCl_2_, 4 Na_2_ATP, 0.4 Na_3_GTP, 5 KCl. This solution was adjusted to pH 7.25 and 295 mOsm. In some experiments K-gluconate was replaced with an equimolar concentration of Cs-gluconate. For experiments requiring intracellular delivery of BAPTA, a modified internal solution was created substituting 10 mM of the primary cation with BAPTA. The liquid junction potential (LJP) was calculated according to the stationary Nernst– Planck equation [25] using LJPcalc (RRID:SCR_025044) and estimated to be 14.8 mV and 15.3 mV for the potassium and cesium based internal solutions, respectively. Results presented are based on calculations that used raw data uncorrected for the LJP.

### 2.4. Protocols

Following establishment of a whole-cell recording, passive membrane properties were evaluated in all neurons using a membrane test protocol that delivered a 200 ms 10 mV hyperpolarizing pulse from a holding potential of −70 mV. Principal neurons were identified by their large soma and regular firing pattern with a maximum firing frequency of < 40 Hz, consistent with previous studies [26,27]. Time course experiments in which access resistance drifted > 30% were also excluded from this study. Whole-cell capacitance was measured using a bidirectional ramp protocol described by Golowasch et al. [28]. Experiments in which membrane test properties were compared before vs. after CCh report mean values from 2.5-4.5 and 8-10 minutes after CCh application relative to the mean value over a 5-minute baseline period before CCh application. Current-clamp experiments evaluating the AHP and/or ADP were performed using a protocol that applied a constant current to hold cells near −70 mV, and a 500 ms depolarizing pulse to cause strong firing (20-40 Hz) determined for each cell individually. AHP and/or ADP measurements were obtained over a 2 second window beginning 100 ms after termination of the depolarizing step and are reported as AUC (area under the curve) relative to the mean voltage over a 1 second period immediately before the depolarizing step. Voltage-clamp experiments evaluating the currents underlying the ADP and/or AHP used a protocol in which neurons were continuously held at −60 mV and stepped to +50 mV for 500 msec. Variations of this protocol recorded multiple sweeps with stimulation duration (0-500 msec in 50 msec steps) or post-stimulation voltage commands of −90 to +20 mV (in 10 mV steps).

### 2.5. Statistics

Population values are presented as mean ± the SEM. Electrophysiological changes before vs. after application of a drug were evaluated for significance in using paired t-tests on raw data, or one-sample t-tests on baseline subtracted data (null hypothesis mean = 0). Curve fitting was performed with OriginPro (Originlab, Northampton, MA).

## 3. Results

### 3.1. Carbachol produces a net excitatory effect on BLA PNs via activation of multiple effectors

Somatic effects of CCh were evaluated in BLA PNs targeted for whole-cell recording in acute brain slices according to their unique morphological and electrophysiological features (see Methods). Passive membrane properties evaluated in control conditions (membrane resistance: 78.71 ± 4.64 MΩ, whole-cell capacitance: 192.31 ± 7.97 pF, n=34) were consistent with previous reports [26]. Holding current at −70 mV was continuously measured as CCh was delivered using a syringe pump to achieve a bath concentration of 10 µM. We found that CCh reliably produced an excitatory shift in holding current (−45.65 ± 3.90 pA, n=34, p=2.59e-13) that plateaued approximately 8 minutes after perfusion began (Figure 1A).

**Figure 1.**
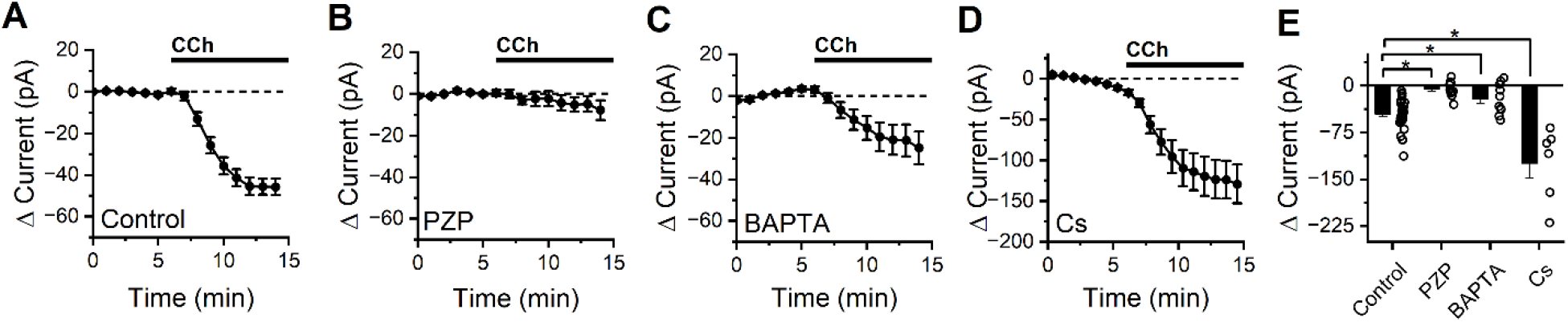
CCh produces an M1/M4 mAChR dependent excitatory current in BLA PNs at subthreshold potentials that is partially blocked by intracellular BAPTA, but not by intracellular cesium. (A) Bath application of CCh causes a negative shift in holding current in BLA PNs voltage clamped at −70 mV, consistent with activation of net excitatory conductances (n=34). (B) Application of the M1/M4 mAChR antagonist pirenzepine (PZP) prevents changes in holding current following CCh exposure (n=10). (C) Intracellular application of BAPTA (10 mM) reduces but does not eliminate the negative shift in holding current produced by CCh (n=6). (D) Intracellular cesium fails to block (in fact increases, see results) overall negative shift in holding current produced by bath application of CCh. (E) Summary data highlighting change in holding current observed in BLA PNs voltage clamped at −70 mV after 10 minutes of CCh exposure, for each condition described in in panels A-D (open circles represent individual cells, bar graphs represent mean ± SE, asterisks indicate p < 0.05).

To gain insight into which mAChR subtype mediates the response to CCh, experiments were repeated in the presence of M1/M4-selective mAChR antagonist pirenzepine (PZP, 1 µM). We found that PZP prevented CCh from causing an excitatory shift in holding current (−5.72 ± 4.03 pA, n=10, p=0.19, Figure 1B) suggesting that the CCh-induced excitatory current was mediated primarily by M1/M4 mAChRs. When the intracellular solution was supplemented with 10 mM BAPTA (a strong calcium chelator), CCh still produced a negative shift in holding current (−21.29 ± 7.57 pA, n=10, p= 0.021, Figure 1C) but it was significantly smaller than the effect observed in control conditions (p=5.31e-03), suggesting that excitation by CCh is at least partially dependent on an effector that is modulated by intracellular calcium. Cesium-based pipette solutions are commonly used to block some types of potassium channels, but when experiments were repeated using a cesium-based intracellular solution we found CCh still produced a large excitatory shift in holding current (−123.82 ± 20.1 pA, n=6, p=3.496e-3, Figure 1D). In fact, this effect was even larger than in observed with a potassium based internal (p=0.02), likely due to the significantly higher basal membrane resistance observed with a Cs-based internal solution (107.8 ± 13.0 MΩ, p=0.02 vs. control conditions).

Interestingly, despite observing an expected strong net excitatory effect of CCh on BLA PNs, we found that CCh reliably produced a biphasic change in membrane resistance (Figure 2A). Specifically, an early decrease in membrane resistance was observed (−9.4 ± 1.1%, p=4.7e-6, n=34, measured 2.5-4.5 minutes after CCh application), followed by an increase in Rm which was apparent by 8-10 minutes after the onset of CCh application (8.3 ± 2.4%, p=1.9e-4, n=34). This finding suggests that CCh may activate at least two distinct types of effectors: one that opens ion channels to lower membrane resistance, and another that closes ion channels to raise it. Consistent with our findings that PZP prevents CCh from changing holding current, we also found that PZP effectively prevents both the observed CCh-mediated decrease and later CCh-mediated increase in Rm (early: −0.66 ± 1.0%, p=0.56, n=10, late: −1.4 ± 1.2%, p=0.48, n=10, Fig. 2B).

**Figure 2.**
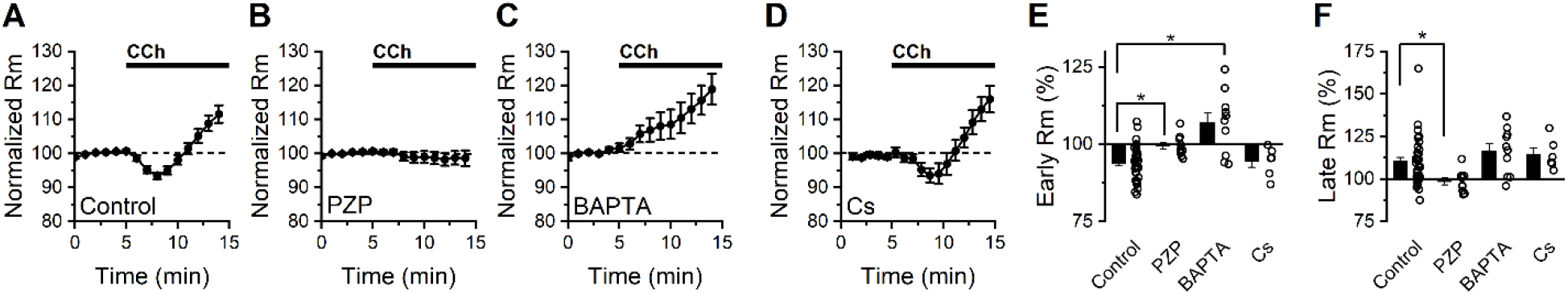
Bath application of CCh has a complex effect of membrane resistance in BLA PNs voltage clamped at −70 mV. (A) Bath application of CCh (10 μM) produces an initial decrease in membrane resistance followed by a later increase (n=34). (B) Continuous application of the M1/M4 mAChR antagonist pirenzepine prevents both the early decrease and late increase in membrane resistance produced by CCh (n=10). (C) Intracellular BAPTA (10 mM) prevents the early CCh-induced decrease in membrane resistance but does not block the later CCh-induced increase in membrane resistance (n=6). (D) Intracellular cesium does not block either the early or late effects of CCh on membrane resistance. E-F) Group comparison of the change in membrane resistance at early and late time points (2.5-4.5 minutes and 8-10 minutes following CCh exposure, respectively).

Next, we noted that when 10 mM BAPTA was included in the internal solution, CCh increased membrane resistance at the early time point (by 7.02 ± 3.20%, p=0.057, n=10), and continued to do so at the later time point (16.5 ± 4.3%, p=0.004, n=10, Fig. 2C). In contrast, when a cesium-based internal solution was used we found that membrane resistance changed similarly to control conditions at both early and late time points (early: −5.59 ± 1.68%, n=6, p=0.039, late: +13.9 ± 3.7%, n=6, p=0.013, Fig. 2D), with no statistically significant difference between cesium and control groups at either time point (p=0.66 early, 0.54 late, vs. control conditions). Collectively, these data in combination with those presented in Fig. 1, suggest that CCh drives depolarization of BLA PNs from near the resting potential in part by activating a cesium-insensitive and calcium-dependent current that drives the membrane potential positive of rest, and also by inhibiting a calcium-independent and cesium-insensitive current that drives the membrane potential negative of rest.

### 3.2. Carbachol enhances an activity dependent afterdepolarization observed in current clamp

CCh has been reported to act on calcium-and voltage-sensitive ion channels to influence excitability of BLA PNs in response to strong firing [21]. To evaluate how mAChR activation influences the AHP and ADP in BLA PNs, membrane potential was evaluated in current-clamp configuration using 500 ms current pulses to elicit strong firing. A continuous current was delivered to each cell to achieve a resting voltage of −70 mV, and the amount of current delivered to each cell to induce strong firing was individually calibrated to maximize consistency across cells (see Methods). Membrane potential after strong firing is reported as the mean voltage for a 2 second window after strong firing relative to the mean voltage before the stimulus. In control conditions, strong firing resulted in an AHP in most cells tested (11 of 14, Figure 3A), with a population mean of −1.42 ± 0.49 mV ^*^ sec (p=0.012, n=14). Following 10 minutes of exposure to 10 µM CCh, strong firing resulted in a large ADP in all cells tested (13.9 ± 0.24 mV ^*^ sec, p=2.99e-4, n=10, Figure 3B), which was significantly more positive than the pre-stimulus baseline (p=1.22e-04). In a subpopulation of cells (2 of 10) the ADP was large enough to reach suprathreshold potentials and cause continuous firing (Figure 3C), a phenomenon that was never observed in the absence of CCh.

**Figure 3.**
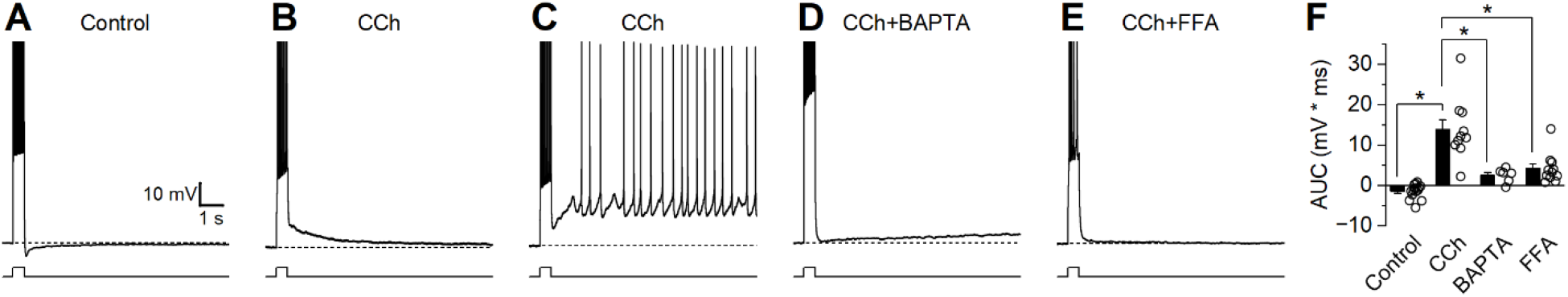
Strong firing in the presence of CCh produces a large ADP. Current-clamp recordings were obtained from neurons held at −70 mV and stimulated with a 500 ms depolarizing current pulse to cause strong firing. (A) Representative trace demonstrating an afterhyperpolarization (AHP) in control conditions. (B) Representative trace demonstrating a large afterdepolarization (ADP) following CCh. (C) Representative trace demonstrating behavior presented by a subset of cells (2 of 10) where CCh resulted in suprathreshold ADPs that caused continuous firing. (D-E) Representative traces demonstrating absence of a large ADP when CCh is applied in the presence of BAPTA or FFA. (F) Group comparison of change in membrane potential after strong firing (Control n=14, CCh n=10, BAPTA n=6, FFA n=11).

Current-clamp experiments evaluating the CCh-induced enhancement of ADP were repeated using modified extracellular or intracellular solutions to gain insight into the signaling pathways that underlie this behavior. When experiments were repeated using a pipette solution containing 10 mM BAPTA, CCh still produced a small ADP (2.55 ± 0.73 mV, p=0.018, n=6, Figure 3D) but it was significantly smaller than the ADP observed without BAPTA (p=2.87e-04) suggesting that much of the ADP is the result of a calcium-dependent intracellular signaling pathway. Similarly, experiments repeated in the presence of I_CAN_ antagonist flufenamic acid (FFA, 100 µM) demonstrated that CCh produces an ADP (4.20 ± 1.12 mV, p=3.75e-3, n=6, Figure 3E) significantly smaller than the ADP observed without FFA (p=4.20e-04). Together these findings suggest that CCh acts on BLA PNs activate an FFA sensitive and calcium-dependent non-selective cation channel, which is primarily responsible for driving the ADP.

### 3.3. Voltage clamp experiments can reveal the detailed kinetics and voltage dependence of the current that underlies the CCh enhanced afterdepolarization

The excitatory current underlying the ADP observed in the presence of CCh (I_ADP_) was further characterized in voltage-clamp configuration using a 500 msec step to +50 mV to simulate strong firing from a −60 mV subthreshold potential. I_ADP_ was measured as area under the curve (AUC) of the voltage-clamp trace for a 3 second period after the depolarizing voltage step relative to the current before the step (see Methods). In the absence of CCh, this protocol allowed the current underlying the AHP (Figure 3A, dashed line) to be measured as an outward current following stimulation (97.4 ± 27.8 pA ^*^ sec, p=0.07, n=3, Figures 4A & 4B). Following 10 minutes of exposure to 10 µM CCh, the depolarizing voltage step resulted in a large inward current, the I_ADP_ (−47.6 ± 11.4 pA ^*^ sec, p=0.05, n=3, Figures 3A & 3B). In order to better isolate and evaluate the I_ADP_, these experiments were repeated with a cesium-based pipette solution, which blocked the potassium channels underlying the AHP that was observed in the absence of CCh. Using this pipette solution, we found a small inward rather than outward current was present in baseline conditions, and the size of this current was significantly increased by bath application of 10 µM CCh (see Fig. 4C for representative example). The I_ADP_ activated by CCh can then be isolated by calculating the difference between the CCh trace and baseline trace, as illustrated in Figure 4D.

**Figure 4.**
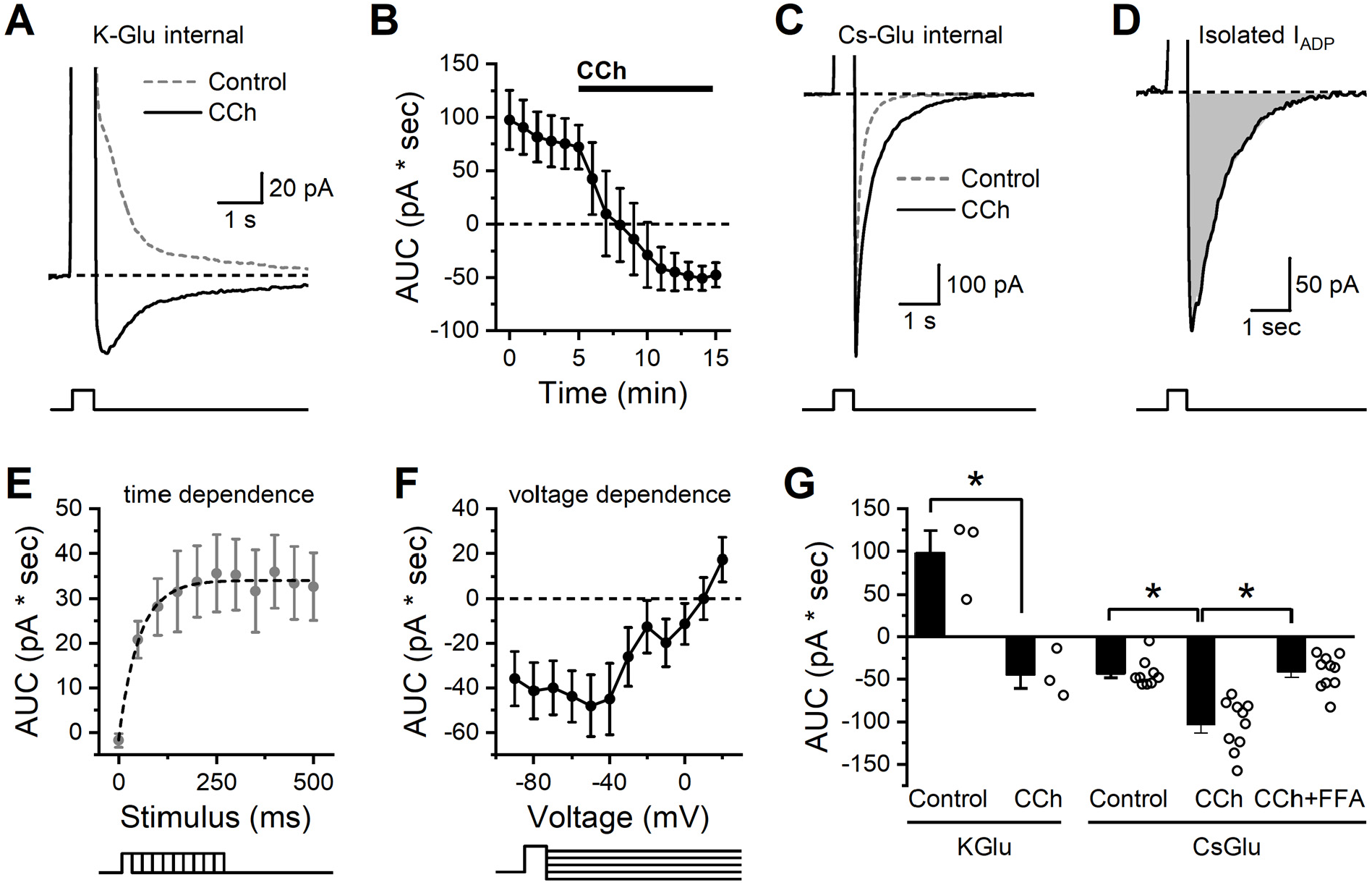
Isolation and characterization of the CCh-activated ADP in BLA PNs. (A) In control conditions a brief but strong depolarization is followed by an outward current (dashed grey line). In the presence of CCh, the same depolarization is followed by an inward current (I_ADP_, solid black line). (B) Repeated measurement of the stimulation-induced current over time further demonstrates that CCh causes a shift from an outward current to an inward current (n=3). (C) The same stimulation using a cesium-based pipette solution prevents the outward current in baseline conditions (dashed grey line) but preserves the ability of CCh to enhance the inward current (I_ADP_, solid black line). (D) The isolated CCh-induced I_ADP_, presented as the difference between CCh and control curves in C. (E) Voltage steps of varying duration were used to reveal that depolarizing steps of 250 msec or greater were required to produce maximal activation of the CCh-induced I_ADP_ (n=11). (F) Voltage dependence of the excitation-activated current demonstrates reversal potential near +10 mV (n=12). (G) Summary data for each cell indicating area of the post stimulus voltage clamp current as observed in both K-gluconate and Cs-gluconate internal solutions after a 250 msec depolarization. Note that the Cs-gluconate internal blocks the outward current observed in control conditions, but CCh still enhances the inward current. Also note that the inward I_ADP_ as isolated here with the Cs-gluconate internal is largely blocked by FFA.

Using this method, we proceeded to evaluate the time and voltage dependence of the isolated CCh-activated I_ADP_. Specifically, experiments that varied the stimulus duration revealed that the maximum I_ADP_ could be produced with a stimulus duration of ~250 msec (Figure 4E). The mean amplitude of the maximum I_ADP_ was 35.3 pA, and the time constant of the activation curve was 52.3 msec. To evaluate voltage dependence, we used a constant stimulus duration of 500 msec, but varied the post-stimulation voltage command. This experiment revealed that the I_ADP_ demonstrated little voltage dependence at subthreshold potentials but had a linear slope at suprathreshold potentials with a reversal potential near +10 mV (Fig. 4F).

To more clearly summarize data across these experiments, in Fig. 4G we present area under the curve of poststimulation currents (recorded at −60 mV after a brief step to +50 mV), both in control conditions and in the presence of CCh. Specifically, the data obtained with a K-gluconate internal (left side of Fig. 4G) further emphasizes that CCh converts the post-depolarization current from outward to inward (n=3, p=0.017), consistent with the current clamp data in Fig. 3 that revealed a CCh-induced shift from an AHP to an ADP. By contrast, in cells filled with Cs-gluconate (right side of Fig. 4G), we observed a post-stimulus inward current in control conditions that was significantly larger when observed in the presence of CCh (n=9,10 for control vs. CCh, p=5.58e-4). Finally, still in cells with Cs-gluconate internal, we noted that if both CCh and FFA were present in the bath, then the post-stimulus current was no larger than observed in control conditions (n=9, 10, respectively, p=0.86), and was also significantly smaller than observed in the presence of CCh without FFA (n=10,10, respectively, p=4.74e-5). These data qualitatively mirror the effects of FFA on post-stimulus responses in current clamp, but importantly represent measurements of a much better isolated cholinergic response. We expect this method to be useful in future studies for making detailed measurements of cholinergic function in a variety of pathological conditions.

## 4. Discussion

This study evaluates the effects of CCh on BLA PNs. In voltage-clamp experiments in which neurons were continuously held at a subthreshold voltage, CCh induced an excitatory shift in holding current with a concomitant biphasic change in membrane resistance (an initial decrease followed by a slower but sustained increase). All these effects of CCh were blocked by 1 μM PZP, suggesting they result from activation of M1/M4 mAChRs. That said, the complex effect on membrane resistance suggests that multiple effectors contribute to the overall excitatory effect of CCh that can be observed in BLA PNs voltage clamped near the resting potential. Indeed, BAPTA partially reduced the overall CCh-induced excitatory response and prevented the early CCh-induced decrease in membrane resistance, without blocking the later increase in membrane resistance. By contrast, cesium failed to reduce the effect of CCh on holding current and did not block either phase of the biphasic effect on membrane resistance. Collectively, these results suggest that CCh can excite BLA PNs in part by opening a calcium-dependent and cesium-insensitive non-selective cation channel, and in part by closing a population of cesium insensitive potassium channels that are open near the resting potential.

CCh was also observed to promote a clear ADP in current clamp, which could sometimes result in sustained firing after a brief stimulus. This effect was similarly prevented by BAPTA, and notably was also blocked by FFA, suggesting that the mechanism of action involves calcium-dependent activation of FFA sensitive non-selective cation channels. We then used voltage clamp protocols to isolate currents after a brief depolarization in both the presence and absence of CCh. In cells filled with a K-gluconate internal, CCh effectively converted a post-depolarization outward current observed in control conditions, to a post-depolarization inward current observed in CCh (that we have referred to as the I_ADP_). These results match the CCh-mediated conversion from an AHP to an ADP after brief depolarization observed in current clamp. In cells filled with cesium gluconate, we noted there was no post-stimulus outward current in control conditions, and that the post-stimulus inward current was both robustly enhanced by CCh, and strongly blocked by FFA. These data strongly suggest that these voltage clamp protocols are enabling us to isolate the current which drives the ADP observed in the presence of CCh in current clamp. Having effectively isolated this current in voltage clamp, we then carefully evaluated both the time and voltage dependence of its activation. In brief, we found a 250 msec depolarization to +50 mV was sufficient to produce near maximal activation of the CCh-induced ADP, and that current carrying the ADP reversed around +10 mV.

Overall, these data are generally in agreement with prior work indicating that BLA PNs have a voltage-and time-dependent potassium current (I_M_) and a calcium-activated K^+^ current (sI_AHP_), both of which are inhibited by CCh, resulting in depolarization [20,22]. Our results also largely align with prior work revealing an FFA sensitive activity dependent current in these neurons [21]. That said, the present study goes to greater lengths to both pharmacologically and mathematically isolate the specific CCh-induced and activity-dependent current that promotes an ADP in current clamp and a post-excitatory inward current in voltage clamp. In broad terms, this current is similar to a calcium- and FFA-sensitive current previously characterized in other brain areas [29–32], although further research may reveal intriguing differences in receptor coupling and in detailed biophysical properties including the voltage and time dependence of activation and of inactivation.

It is worth highlighting that within the BLA there are clearly multiple types of both ionotropic and metabotropic cholinergic receptors, a number of which can be activated by CCh [21,33–35]. In our view the clear effectiveness of PZP against multiple effects of CCh observed in this study in BLA PNs voltage clamped at −60 mV leads us to conclude that the vast majority of the responses apparent throughout this study are very likely mediated by activation of M1 and/or M4 AChRs [36]. Nevertheless, it seems likely nicotinic AChRs and or M2 containing mAChRs impact neuronal excitability in BLA in other contexts. Interestingly, Yajeya et al. [37] reported that BLA PNs also have a calcium-independent nonselective cation conductance that is activated by CCh. If this current were present in our experiments, it could plausibly contribute to the change in holding current observed in the presence of BAPTA in Figure 1C. Similarly, although FFA acts on a few different ion channels [38], collectively our data strongly suggest that an FFA-sensitive and calcium-activated non-selective cation channel is being engaged by direct excitation of BLA PNs in the presence of CCh. It would likely be interesting for further studies to determine whether transient receptor potential (TRP) cation channels contribute directly to the CCh-induced ADP observed in these neurons. Interestingly, Meis et al. [11] provide compelling evidence for the presence of a nonselective cation channel in mouse BLA projection neurons that is activated downstream of CCK receptors, is sensitive to TRP channel blockers, and is not dependent on increases in intracellular calcium.

Overall, this work provides additional insight into the multiple cholinergic effectors that modulate the intrinsic excitability of BLA PNs, and particularly demonstrates methods that can be used to isolate the current underlying classic ADPs and plateau potentials, so that their biophysical properties can be more precisely measured, and ultimately compared across healthy and pathophysiological conditions.

## 7. Author Contributions

Conceptualization: T.J.S., S.W.H., J.L.B., B.S. and C.J.F.; Formal analysis: T.J.S. and S.W.H.; Investigation: T.J.S. and S.W.H.; Project administration: C.J.F.; Software: S.W.H. and C.J.F.; Supervision: B.S. and C.J.F.; Visualization: T.J.S. and S.W.H.; Writing – original draft: T.J.S. and S.W.H.; Writing - review & editing: J.L.B., B.S., and C.J.F.

